# Transcriptome characterization of the common cockle (*Cerastoderma edule*) after exposure to a *Marteilia cochillia* outbreak

**DOI:** 10.1101/2022.10.18.512677

**Authors:** B. G. Pardo, C. Fernández, M. Pampín, A. Blanco, D. Iglesias, A. Cao, M.J. Carballal, A. Villalba, P. Martínez

**Affiliations:** UNIVERSIDADE DE SANTIAGO DE COMPOSTELA; Centro de Investigacions Marinas, Conselleria do Mar, Xunta de Galicia; Universidade de Santiago de Compostela

**Keywords:** marteiliosis, RNA-seq, differentially expressed genes, resilience, immune response

## Abstract

The edible cockle (*Cerastoderma edule*) is a widely cultivated bivalve with relevant ecological value roles and high value for shellfisheries in different European regions. The emergence of new threats, such as the parasite *Marteilia cochillia*, has impaired the production and ecosystem of shellfish beds where the parasite was detected. Knowledge of the molecular mechanisms involved in cockle immune response to this parasite is essential to devise strategies for its control. With this aim, a transcriptomic study of the digestive gland (target organ of the parasite) and the whole cockle meat in response to *M. cochillia* infection was carried out in heavily impacted area in the Northwest of Spain (Lombos do Ulla, Ria de Arousa). A total of 2079 million raw RNA-seq reads were obtained after filtering and used for annotation of 9049 genes following a conservative bioinformatic pipeline using the chromosome-level cockle genome as reference. Gene expression analysis identified a total of 973 consistent differentially expressed genes (DEGs) between comparisons across a temporal series involving cockles with different degrees of infection. DEGs increased with the level of infection within each temporal sample, but the higher DEGs number were detected when comparing temporal samples. Enrichment analysis of DEGs showed an increased expression of molecular functions related to hydrolase, peptidase activity, carbohydrate binding and active transmembrane transporter activity; cellular components such as extracellular matrix and extracellular regions; and a few biological functions associated with immunity and defence response. This information will be valuable for further studies focused on DEGs and associated SNP markers to develop reliant cockle strains to marteiliosis.

## Introduction

The common cockle *Cerastoderma edule* (Linnaeus, 1785) is a shellfish species with a wide geographical distribution throughout the European coast from the Barents Sea and the Baltic Sea to the Iberian Peninsula, and southwards along the coast of West Africa to Senegal (Tebble, 1976). Cockle is one of the most abundant mollusc species in European bays and estuaries where population densities of ~10,000 per m^2^ have been recorded (Tyler-Walters, 2007). In those areas where it lives, the common cockle contributes significantly to the coastal ecosystems and plays an important socioeconomic role. The most prominent service provided by cockles to human communities is food production, but many other important services are provided, such as the use of shells for ornaments, poultry grit and construction (Kelley, 2009; Morris et al., 2018; van der Schatte Olivier et al., 2020; Jackson-Bué et al., 2022). Also, its role in coastal ecosystems has been highlighted since cockles contribute very effectively at increasing the productivity of sedimentary habitats, and directly provide a food source for predators, thereby supporting the biodiversity and productivity of a wide range of other species (Carss et al., 2020; Dairain et al., 2020).

Cockles are commercially important in the UK, Ireland, Netherlands, Portugal, Spain, and France. In recent decades, cockle populations have declined across Europe, with production dropping from 108,000 tons in 1987 to 26,100 tons in 2019 (FAO, 2021). Common cockle decline has been attributed to different factors according to locations, such as predation, diseases, climatic events, pollution, poor recruitment and over-fishing and parasitic infections (Ducrotoy et al., 1991; Wolff, 2005; Thieltges, 2006; Longshaw and Malham, 2013). Cockles host a wide variety of normally innocuous viruses, bacteria, fungi and protistans, but they can become problematic in certain environmental scenarios. In Galicia (NW Spain), unprecedented mass mortalities attributed to marteiliosis (infection caused by the protistan parasite *Marteilia cochillia*), led to cockle fishery collapse in Ria de Arousa in 2012 (Villalba et al., 2014). From this area, the parasite expanded southwards to beds in the Ria de Pontevedra and Ria de Vigo in 2013 and 2014, respectively (Iglesias et al., 2015, 2017, 2019). The absence of marteiliosis outbreaks in the cockle beds northwards, on the other hand, has allowed Ria de Muros-Noia to be the most productive cockle area in Galicia (Iglesias et al., 2015).

Molluscs possess a powerful innate defense system but lack the lymphocyte-based system that characterizes vertebrate immunity (Schultz and Adema, 2017). Therefore, standard treatments for infectious disease control in vertebrates cannot be applied because they are mainly cultured in open areas and true vaccines cannot be developed because molluscs lack immune memory, although adaptive immune mechanisms have recently been claimed (Pradeu and Du Pasquier, 2018; Yao et al., 2021). Their immune response relies on innate cellular and humoral mechanisms, both operating in coordination to provide protection against pathogens. The innate immune system of bivalves can be summarized in three main steps: (i) the recognition of molecular motifs by soluble compounds and cellular receptors associated with microorganisms or endogenous molecules secreted by damaged tissues, (ii) the activation of different signaling pathways, and (iii) the production of molecular effectors involved in host and cellular defense responses (Allam and Raftos, 2015; Wang et al., 2018; Watson et al., 2022). The better the comprehension of the interaction between the immune system of cockles and the molecular mechanisms of pathogen infectivity, the more efficient the fight against diseases.

In this context, developing large-scale genomic resources is a priority to facilitate the implementation of new technological approaches that can aid in the identification of signalling networks controlling disease resistance (Gómez-Chiarri et al., 2015). Genomic technologies are advancing rapidly and becoming more affordable and accessible (Potts et al., 2021). RNA-seq is an efficient high-throughput method that has provided a wealth of genomic information for non-model species. Transcriptome sequencing has several advantages including the identification of genes and associated markers (SNP: Single Nucleotide Polymorphism), good repeatability and quantitative criteria, and further, they are cost-effective (Nagalakshmi et al., 2008; Houston et al., 2020). Transcriptomic information can be used to quantify gene expression profiles, characterize functional-related genes, and provide insights into the biological issues addressed. Understanding the molecular basis of the differences responsible for being susceptible or resistant to a particular disease and identifying molecular markers of resistance can help to fight against mollusc diseases. Recently, research effort has been focused on the identification of molecular markers of disease resistance, through transcriptomics (He et al., 2012; Meistertzheim et al., 2014; Nikapitiya et al., 2014; Nie et al., 2015; Wang et al., 2016; Gutierrez et al., 2018, 2020; La Peyre et al., 2019; De Lorgeril et al., 2020; Farhat et al., 2020; Proestou and Sullivan, 2020; Hasanuzzaman et al., 2020), proteomics (Simonian et al., 2009; Fernández-Boo et al., 2016; de la Ballina et al., 2018; Vaibhav et al., 2018; Smits et al., 2020; Leprêtre et al., 2021) and population genomics approaches (Vera et al., 2019; Sambade et al., 2022). This information could be eventually used for marker-assisted selection (MAS) programs to increase resilience against pathogens.

In this study, we used an RNA-seq approach to characterize the common cockle transcriptome after exposure to *M. cochillia* along a temporal series and different degrees of infection. We focused mainly on the digestive gland, the target organ of this parasite, but also obtained information from the whole meat at different life stages. Results provided a baseline for further understanding the transcriptomic mechanisms involved in immune response at the molecular level with the aim of detecting differentially expressed genes (DEGs) and associated SNPs that could be further used for MAS by industry or administration to recover cockle production using marteiliosis-resistant strains.

## Materials and methods

### Experimental design and sample collection

Previous research on cockle-marteiliosis dynamics in the shellfish bed of Lombos do Ulla (Ria de Arousa) (42° 37,757′ N, 8° 46,521′ W) showed an epidemic annual pattern: marteiliosis outbreaks start in summer or early autumn and cause massive mortality and depletion of nearly the whole recruited cohort by late winter to early spring, before achieving the minimum market size (Iglesias et al. submitted). Accordingly, we devised our experiment to understand the genetic response to marteiliosis across one year outbreak from July 2018 to July 2019. The cockle digestive gland is the target organ for *M. cochillia* infection and proliferation, so we mainly focused on this organ for transcriptomic comparisons over time and across the different levels of infection, although for some samples it was unavoidable dissecting the whole meat, either by their small size or by experimental design issues.

**Figure 1.**
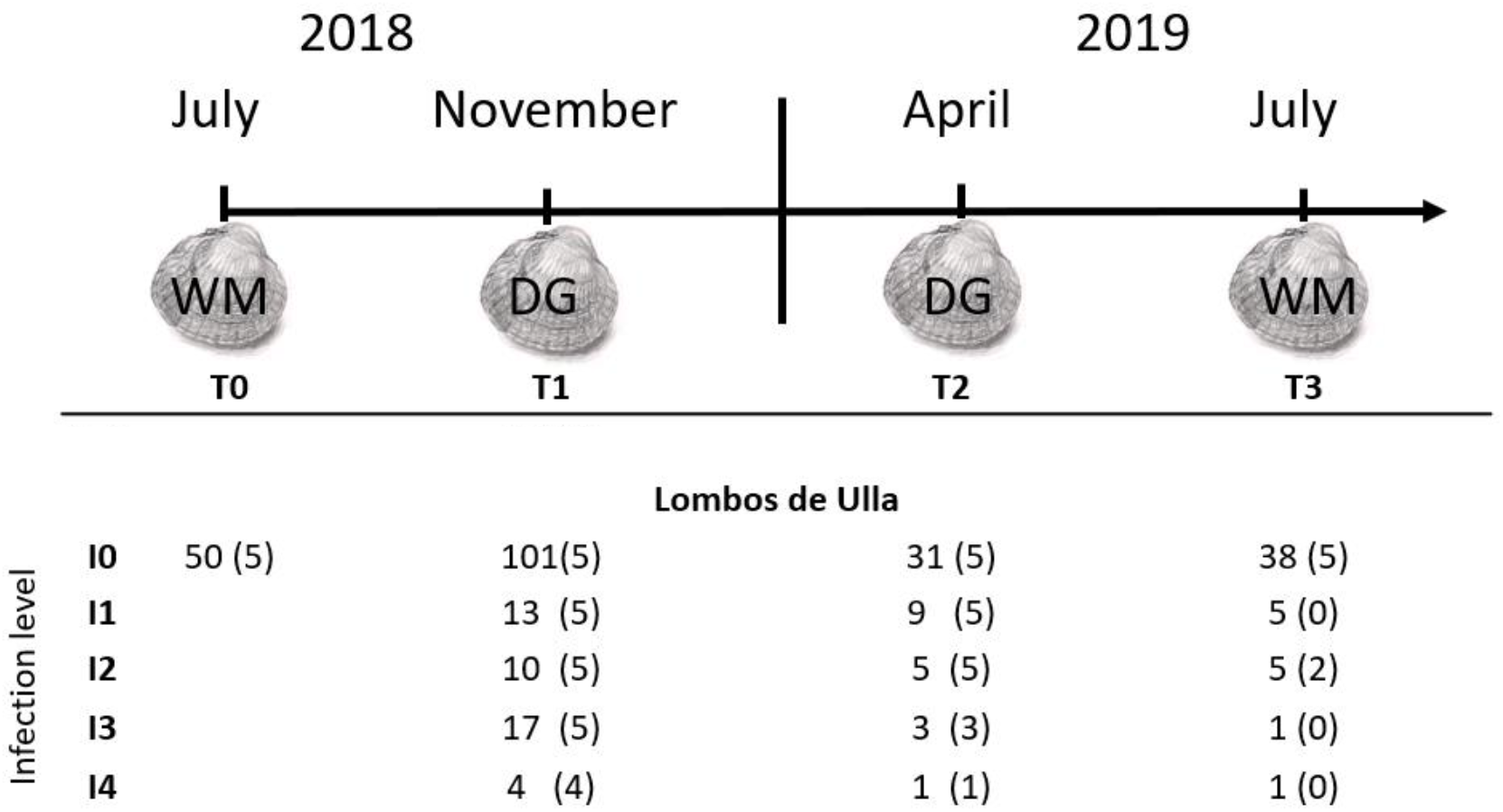
Sampling of *Cerastodermma edule* in Lombos do Ulla for transcriptomic analyses. WM: Sample of whole cockle meat; DG: Sample of digestive gland; I0 to I4: increasing levels of infection. In parentheses the number of samples used for RNA-seq analysis.

The experiment started when a newly recruited cohort was settled in Lombos do Ulla in July 2018 and finished one year later when most of the population was nearly depleted. Samples of the digestive gland or the whole meat were taken to the laboratory for transcriptomic analysis and for histopathological evaluation to categorize the level of infection. Four sampling times were accomplished along the year evaluated: T0, before the first detection of *M. cochillia (July 2018)*; T1 and T2, as two intermediate times in November 2018 and April 2019, respectively; and T3, the last time in July 2019. It was intended a representation of five individuals per time and level of infection, although it was not always possible in the samples collected (Fig. 1). Considering the small size of T0 cockles, a sample of five cockles from the shellfish bed of Noia, located in the northern neighbor Ria de Muros-Noia, where marteiliosis outbreaks were never reported, was collected to have a reference digestive gland from non-infected individuals to be used as uninfected controls for RNA-seq analysis. In addition, due to the low survival at the end of the experiment (T3), the whole meat of each individual was divided in two parts to complete all necessary samples for histopathological and transcriptomic analysis, so in this case, half of the whole meat was used for RNA extraction.

### Histopathological analysis

To categorize the marteilioisis level of infection, we used the procedure described by Villalba et al. (2014) and Iglesias et al. (submitted). Briefly, the cockles collected were carried to the laboratory and kept in a tank with open seawater flow for 24 h to allow the elimination of gut content. Then, a transversal section (about 5 mm thick) of soft tissues, including part of the digestive gland, was fixed for staining with Harris’ haematoxylin and eosin (Howard et al., 2004) to be examined under light microscopy to diagnose *M. cochillia* infection. Cockles were classified as follows: I0, non-infected; I1; early infection stage; I2, moderate infection stage; I3, heavy infection stage; and I4, final infection stage, where the digestive gland appeared highly degraded due to parasite sporulation stages that filled the intestinal lumen (de Montaudouin et al., 2021; Iglesias et al. submitted) (Fig. 1).

### RNA extraction and RNA-seq

Digestive gland or whole meat samples were stored at −20°C in RNAlater for one night at 4°C until processing. Total RNA was individually extracted using the RNeasy Mini Kit (Qiagen, Hilden, Germany) with slight modifications, including two additional RPE washes and the optional DNase digestion. RNA purity was determined with a NanoDrop® ND-1000 spectrophotometer (NanoDrop® Technologies Inc.). RNA integrity was assessed with the RNA nano 6000 assay kit of the Agilent Bioanalyzer 2100 system (Agilent Technologies, CA, USA). In molluscs, the calculation of RNA integrity number (RIN) is usually affected by the co-migration of 28S and 18S rRNA fragments (Barcia, 1997), so, a single peak with a flat baseline was used to assess the RNA integrity.

A total of 60 good RNA quality (RIN > 8.1) samples, including five non-exposed individuals from Noia (digestive gland control) and 55 individuals from Lombos do Ulla (5 non-exposed and 50 exposed) collected at four times (T0, T1, T2, T3) and representing different infection levels (from I0 to I4) were delivered for RNA-seq (Fig. 1). Samples were barcoded and prepared for sequencing by Novogene (UK) Company Limited on Illumina NovaSeq S2 as 150 bp paired-end reads.

### RNA-seq data processing and functional annotation

The whole transcriptomic information of the samples collected was used to reconstruct the cockle’s transcriptome, mostly constituted of digestive gland information, but also including the whole meat (T0 and T4; Fig. 1). The quality of the sequencing output was assessed with FastQC v.0.11.7 (https://www.bioinformatics.babraham.ac.uk/projects/fastqc/). Quality filtering and removal of residual adaptor sequences were performed using Trimmomatic v.0.39 (Bolger et al., 2014). Specifically, i) residual Illumina-specific adaptors were clipped from the reads; ii) the read removed, if a sliding window average Phred score over five bases was < 20; and iii) only reads where both pair-ends were longer than 50 bp post-filtering were retained. Filtered reads were aligned against the cockle genome (Bruzos et al., 2022) using a custom script with SAMtools v1.9 (Li et al., 2009) and STAR v2.7.0e (Dobin et al., 2013). Bam files from aligned reads were merged with SAMtools and a cockle transcriptome draft was finally obtained. Gene detection and annotation were accomplished with StringTie v1.3.5 (Pertea et al., 2015) and BLASTx (Evalue < 10^−5^; (Altschul et al., 1990). A pipeline was designed to avoid redundancy and obtain the most reliable transcriptome to be used for further analyses (Fig. 2): i) we collapsed those overlapping genes with identical annotation; and ii) genes without annotation, uncharacterized annotation or in small genome scaffolds not anchored to chromosomes were discarded. GO terms for each sequence were obtained by blasting it against the uniref90 database, taking the best homologous sequence, and using its name to extract the GO terms from the corresponding web page at the web portal uniprot.org using a custom Perl script. Transcriptome functional characterization was made using WEGO (Ye et al., 2006, 2018) and Singular Enrichment Analysis (SEA) was performed in the website platform AgriGO v2.0 (Tian et al., 2017) using a 5% false discovery rate (FDR).

### Identification of differentially expressed genes (DEGs) and markers across the infection process

Transcript abundance (expressed as TPM: transcripts per kb of the gene per million reads) was obtained by pseudo-aligning the filtered RNAseq reads from each individual against the cockle genome reference with Kallisto 0.46.1 (Bray et al., 2016) using 100 bootstrap samples (-b100). Identification of differentially expressed genes (DEGs) was performed with Sleuth (Pimentel et al., 2017) using a Wald Test (WT) and statistical significance was established with a Benjamini-Hochberg multiple testing corrected q-value ≤ 0.05.

**Figure 2.**
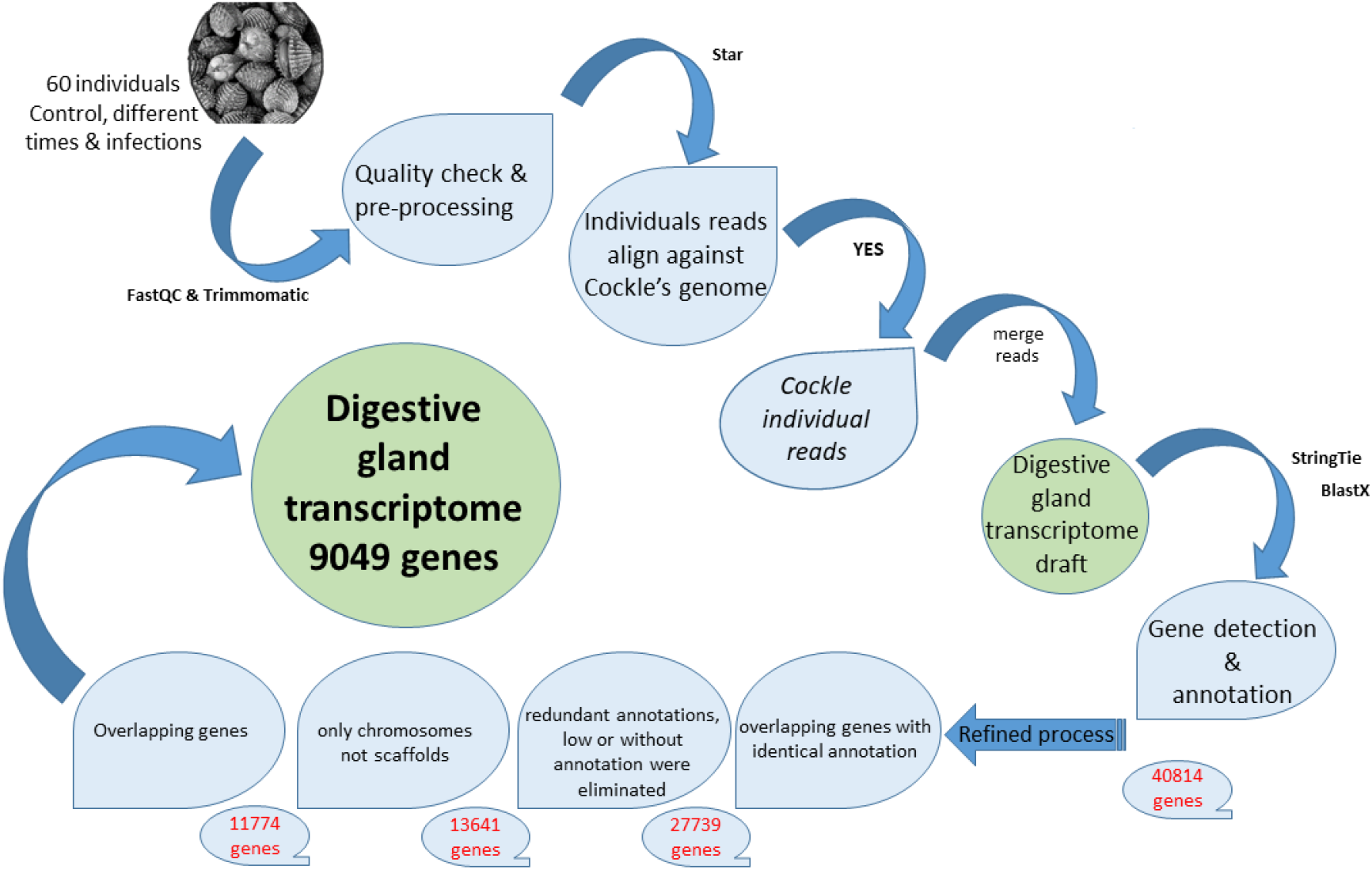
Bioinformatic pipeline for *C. edule* transcriptome characterization.

### DEGs across time and level of infection

Transcriptomes were compared across conditions considering the time and level of infection, always using the same tissue (digestive gland or whole meat) (Fig. 3). Two temporal pair-wise comparisons were accomplished: i) T1 *vs* T2 vs Noia using digestive gland; and ii) T0 *vs* T3 using the whole meat. Also, different infection levels (I0, I1, I2, I3, I4) were pairwise compared within each time (T1, T2) taking I0 as reference (Fig 3).

**Figure 3.**
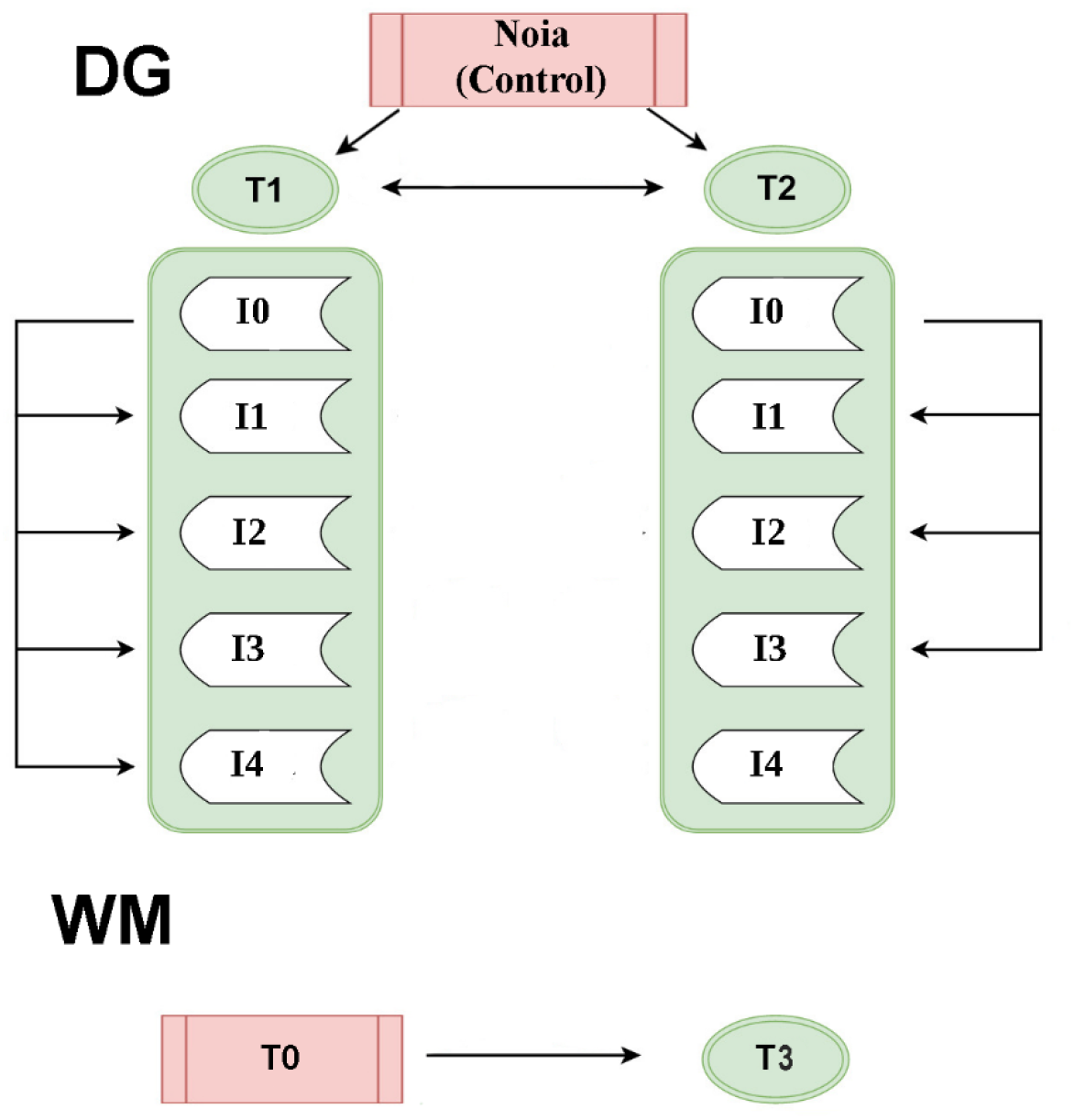
Comparisons performed to identify differentially expressed genes in the *Cerastoderma edule* digestive gland (DG: T1, T2) and whole meat (WM: T0, T3) with different infection degrees of a *Marteilia cochillia* (I0 to I4, from low to heavy infection) from the outbreak occurred in Lombos do Ulla in 2018-2019.

## Results and Discussion

*Marteilia cochillia* was detected for the first time in the Mediterranean coastal waters of Spain in 2008 during investigations on *C. edule* mortalities (Carrasco et al., 2011, 2013), despite the typical cockle species in the Mediterranean coasts is the lagoon cockle (*C. glaucum;* Malham et al., 2012). Four years later, it was identified as the causative agent of cockle decline on the Atlantic coast of Spain (Villalba et al., 2014). Recently, new mortality events associated with *M. cochillia* have been reported in both Mediterranean and Atlantic locations (Carrasco et al., 2013; Iglesias et al., 2017). The spreading of the parasite in Galician rias has contributed to the collapse of cockle fisheries, causing the extinction of every newly recruited cockle cohort before reaching market size (Iglesias et al., 2017, 2019), highlighting its extreme pathogenicity. In this study, we tackled for the first time the characterization of the transcriptome response of common cockle to this parasite which is devastating its production in the Southern Galician rias (NW Spain) as a first step for identifying candidate genes and markers to control this parasitosis.

### RNA sequencing output

A total of 2079 million 150 bp paired-end raw reads were generated, representing on average 34.6 million reads for each of the 60 samples analyzed, 1681 million from the digestive gland and 398 million from the whole meat samples (Table 1). After filtering, 1991 million total reads were retained, 1609 million from the digestive gland and 381 million from whole meat. Among the 33 million pair-end filtered reads per sample, 28.8 million were consistently aligned to the chromosome-level assembled cockle genome (Bruzos et al., 2022) by retaining only the single best hit when more than one position matched.

**Table 1.**
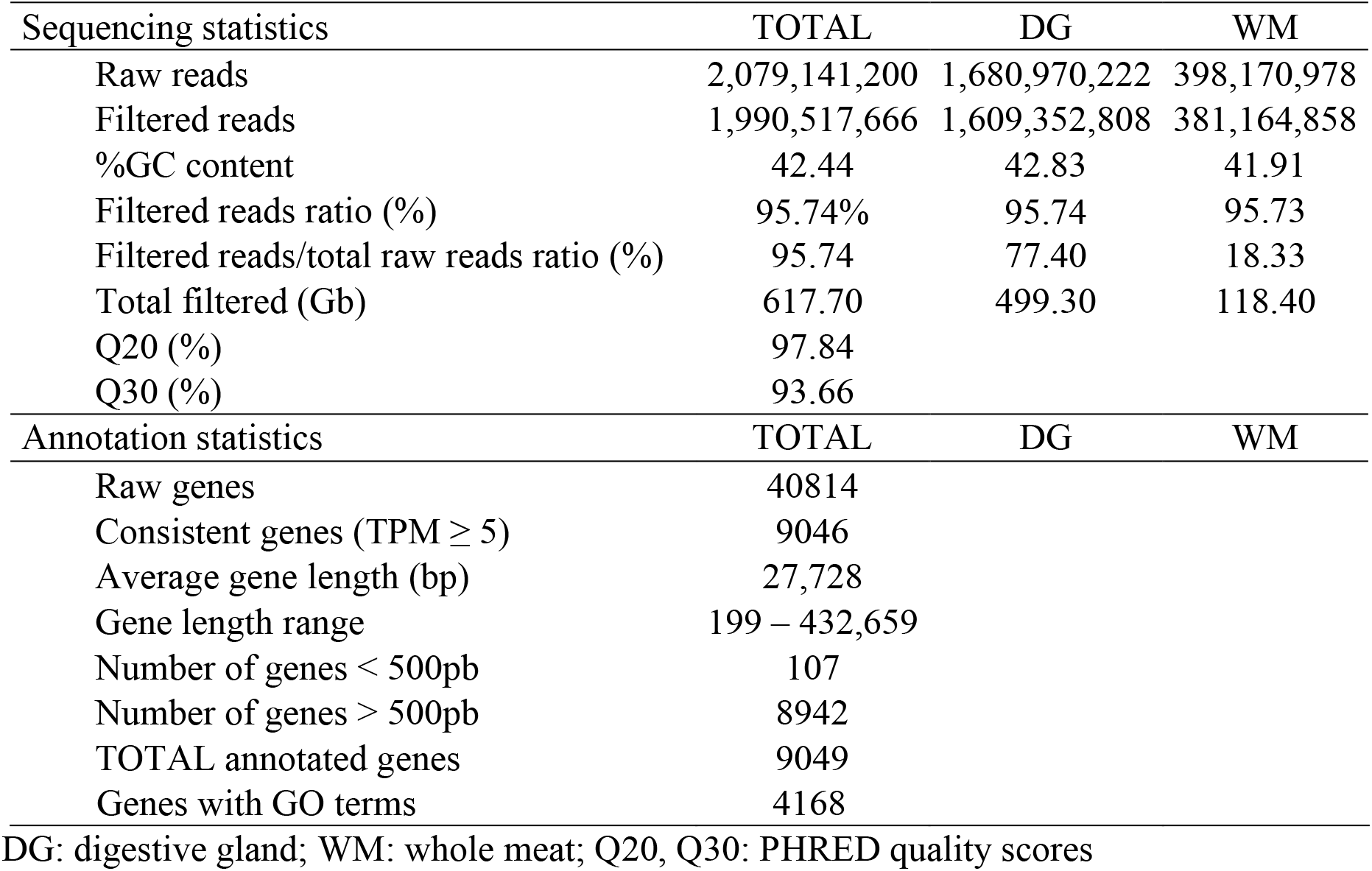
Summary of sequencing and assembly transcriptome data of *Cerastoderma edule* from a *Marteilia cochillia* outbreak in Lombos do Ulla.

**Figure 4.**
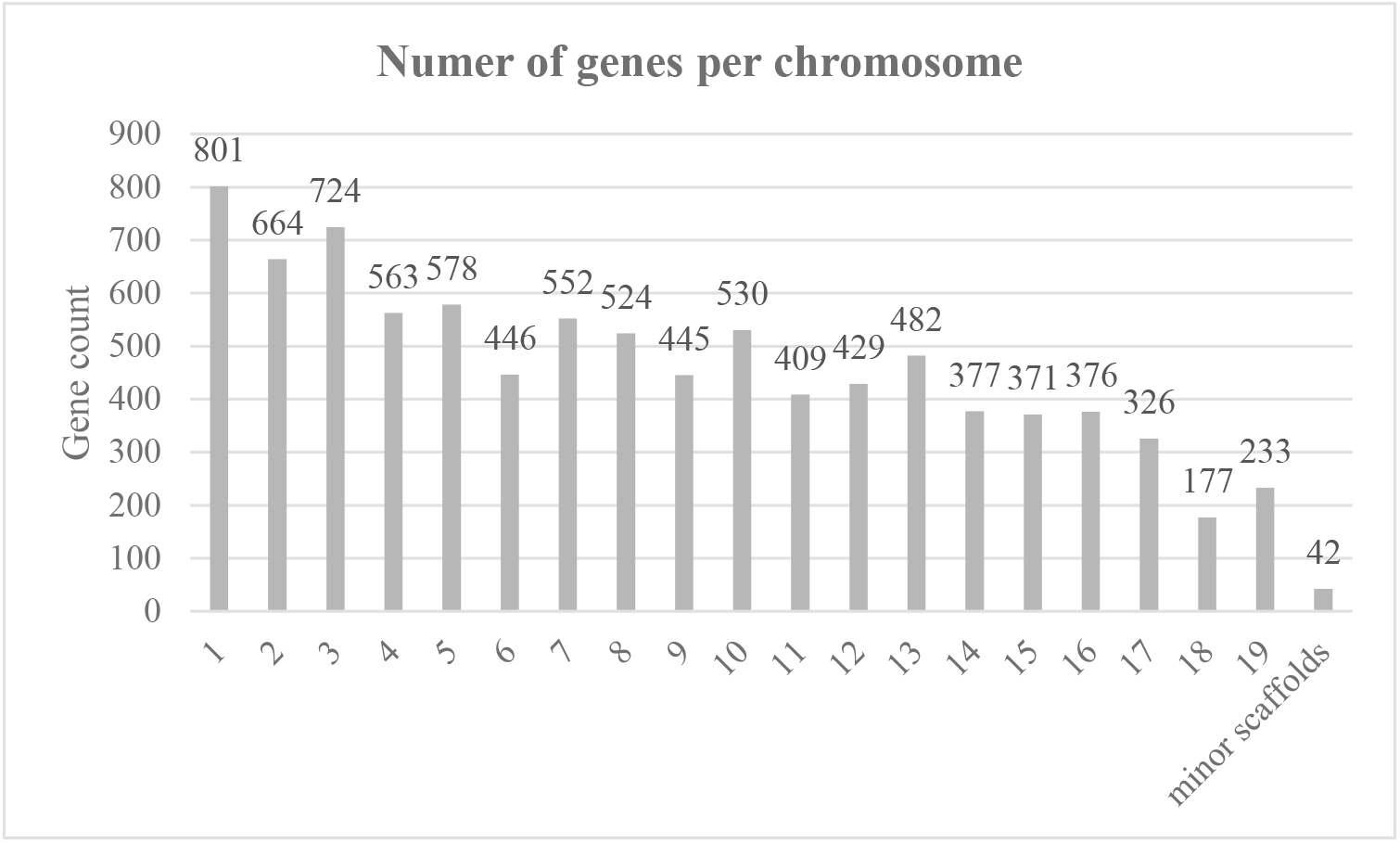
Frequency histogram of annotated gene number per chromosome of *Cerastoderma edule* transcriptome.

The common cockle transcriptome, particularly enriched on the digestive gland, was constructed taking as reference the cockle genome assembly and using a conservative bioinformatic pipeline (Fig. 2): i) overlapping genes with the same annotation were collapsed to a single gene and the length of the overlapping region was considered as their full length; ii) those genes with no annotation were discarded; and iii) only those genes with ≥ 5 TPM in at least one sample were retained. Following this approach, from the initial 48,214 annotated genes, a total of 9046 consistent genes constituted the cockle’s transcriptome according to our filtering criteria (Table S1). Only three putative genes with ≤ 3 reads were discarded. A total of 7421 genes showed significant expression (≥ 5 TPM) in more than 50% of the samples and 5689 genes showed high expression (≥ 1000 TPM) considering all samples (Table 2).

**Table 2.**
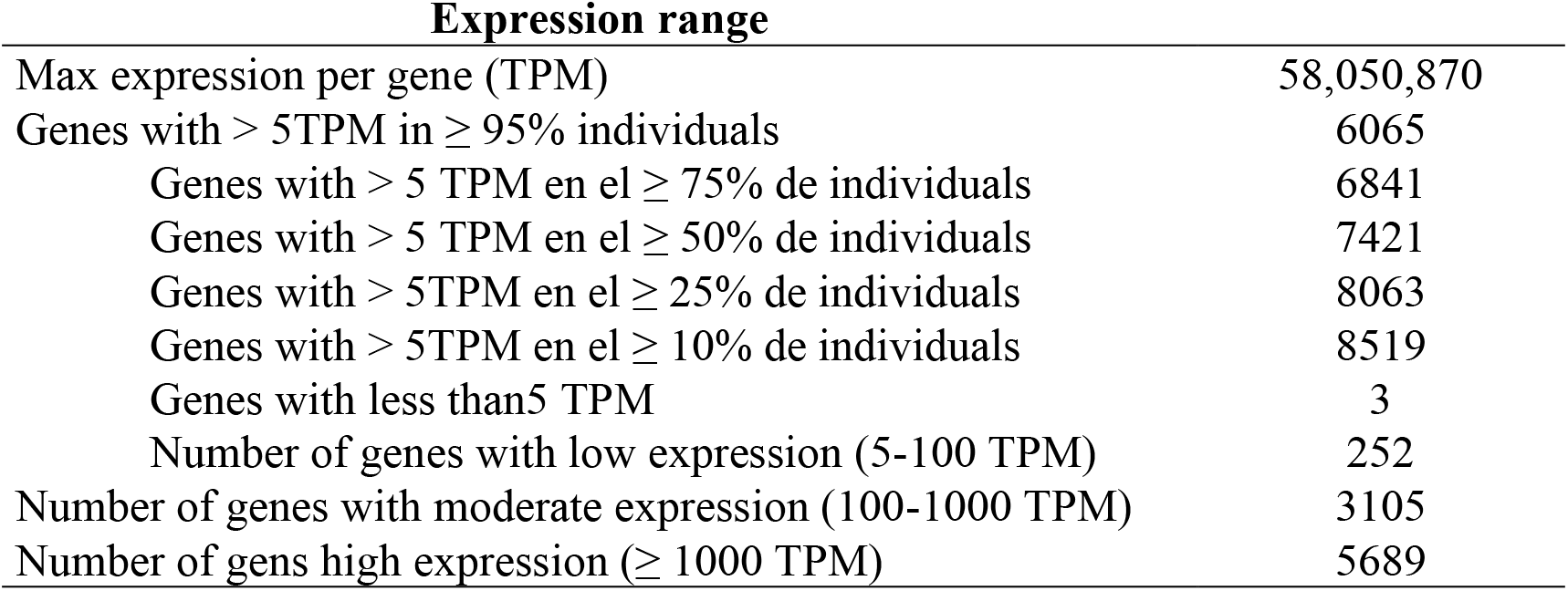
Gene expression range in the *Cerastoderma edule* transcriptome.

| **Expression range** |  |
| --- | --- |
| Max expression per gene (TPM) | 58,050,870 |
| Genes with > 5TPM in $\geq 95\%$ individuals | 6065 |
| Genes with > 5 TPM en el $\geq 75\%$ de individuals | 6841 |
| Genes with > 5 TPM en el $\geq 50\%$ de individuals | 7421 |
| Genes with > 5TPM en el $\geq 25\%$ de individuals | 8063 |
| Genes with > 5TPM en el $\geq 10\%$ de individuals | 8519 |
| Genes with less than 5 TPM | 3 |
| Number of genes with low expression (5-100 TPM) | 252 |
| Number of genes with moderate expression (100-1000 TPM) | 3105 |
| Number of gens high expression ($\geq 1000$ TPM) | 5689 |

The distribution of genes per chromosome (C) was roughly in accordance with its length, ranging from 177 in C18 to 801 in C1, with 42 genes being localized in minor unanchored scaffolds (Fig. 4). Using Kallisto read count, we extracted the 20 most expressed genes that resulted to be homologous to other molluscs, mostly bivalves, in the nr database (Table 3; Table S2). These genes were mainly associated with ribosomal constitution and immune response, indicating an active protein synthesis and response to marteiliosis infection.

**Table 3.** List of the 20 most expressed genes in *Cerastoderma edule* transcriptome normalized by TPM. Blastx search in non-redundant protein sequences (nr).

| **Gene annotation** | **Species best hit** | **Accession code** | **E value** | **TPM** |
| --- | --- | --- | --- | --- |
| Cytochrome b | *C. edule* | YP_009420861.1 | 4.00E-84 | 3483052.90 |
| Uncharacterized protein LOC123527942 | *M. mercenaria* | XP_045163626.1 | 1.00E-23 | 982498.14 |
| Plasma kallikrein-like | *M. mercenaria* | XP_045213758.1 | 8.00E-145 | 923517.34 |
| Integrin beta | *Solen grandis* | ATB53129.1 | 5.00E-45 | 547607.17 |
| Hypothetical protein BaRGS_027580 | *Batillaria attramentaria* | KAG5691076.1 | 0.00E+00 | 534569.05 |
| 40S ribosomal protein S2 | *M. mercenaria* | KAH3866819.1 | 2.00E-155 | 470436.70 |
| Core histone H2A/H2B/H3/H4 | *Dictyocaulus viviparus* | KJH53109.1 | 8.00E-90 | 463564.57 |
| Zinc metalloproteinase nas-7-like | *M. mercenaria* | XP_045212251.1 | 6.00E-136 | 404317.11 |
| 40S ribosomal protein SA-like | *M. mercenaria* | XP_045213159.1 | 1.00E-160 | 381382.45 |
| 60S ribosomal protein L27-like | *M. mercenaria* | XP_045169850.1 | 1.00E-61 | 377310.89 |
| Hypothetical protein DPMN_010003 | *Dreissena polymorpha* | KAH3886003.1 | 2.00E-93 | 375811.82 |
| 60S ribosomal protein L19-like | *M. mercenaria* | XP_045181329.1 | 8.81E-01 | 353062.74 |
| Actin, cytoplasmic | *M. mercenaria* | XP_045158456.1 | 8.00E-92 | 342245.76 |
| Hypothetical protein DPMN_121632 | *Dreissena polymorpha* | KAH3819888.1 | 4.00E-62 | 331076.26 |
| Ribosomal protein S25 | *Lepidochitona cinerea* | ACR24971.1 | 1.00E-38 | 330242.42 |
| Slit homolog 1 protein-like | *M. mercenaria* | XP_045199480.1 | 0.00E+00 | 329901.52 |
| 40S ribosomal protein S13 | *M. mercenaria* | XP_045176593.1 | 1.00E-95 | 321114.58 |
| 40S ribosomal protein S9 | *M. mercenaria* | XP_045211562.1 | 1.00E-111 | 320790.15 |
| Adenosylhomocysteinase A-like | *M. mercenaria* | XP_045193115.1 | 0.00E+00 | 300424.49 |
| 60S ribosomal protein L18a-like | *M. mercenaria* | XP_045165720.1 | 9.00E-104 | 293144.62 |

One or more GO terms were retrieved from the best hit for 4186 of the overall 9046 genes (46.3%), of which 2393 corresponded to cellular component (CC), 3351 to molecular function (MF) and 1932 to biological process (BP) categories (Fig. 5). The most descriptive category, BP, showed a wide variety of functions, being those related to cellular, metabolic, and biological regulatory processes the most relevant. Other important functions such as response to stimuli and signalling showed also a relevant representation. As for MF, most of the genes were related to binding and catalytic activity, while for CC functions were associated with cell parts, membrane and membrane parts and organelle (Fig. 5; Fig. S1). Overall, the annotation and functional classification of *C. edule* transcripts provided a valuable dataset for the identification of functional genes and pathways as well as for further genome research and analysis in this species.

**Figure 5.**
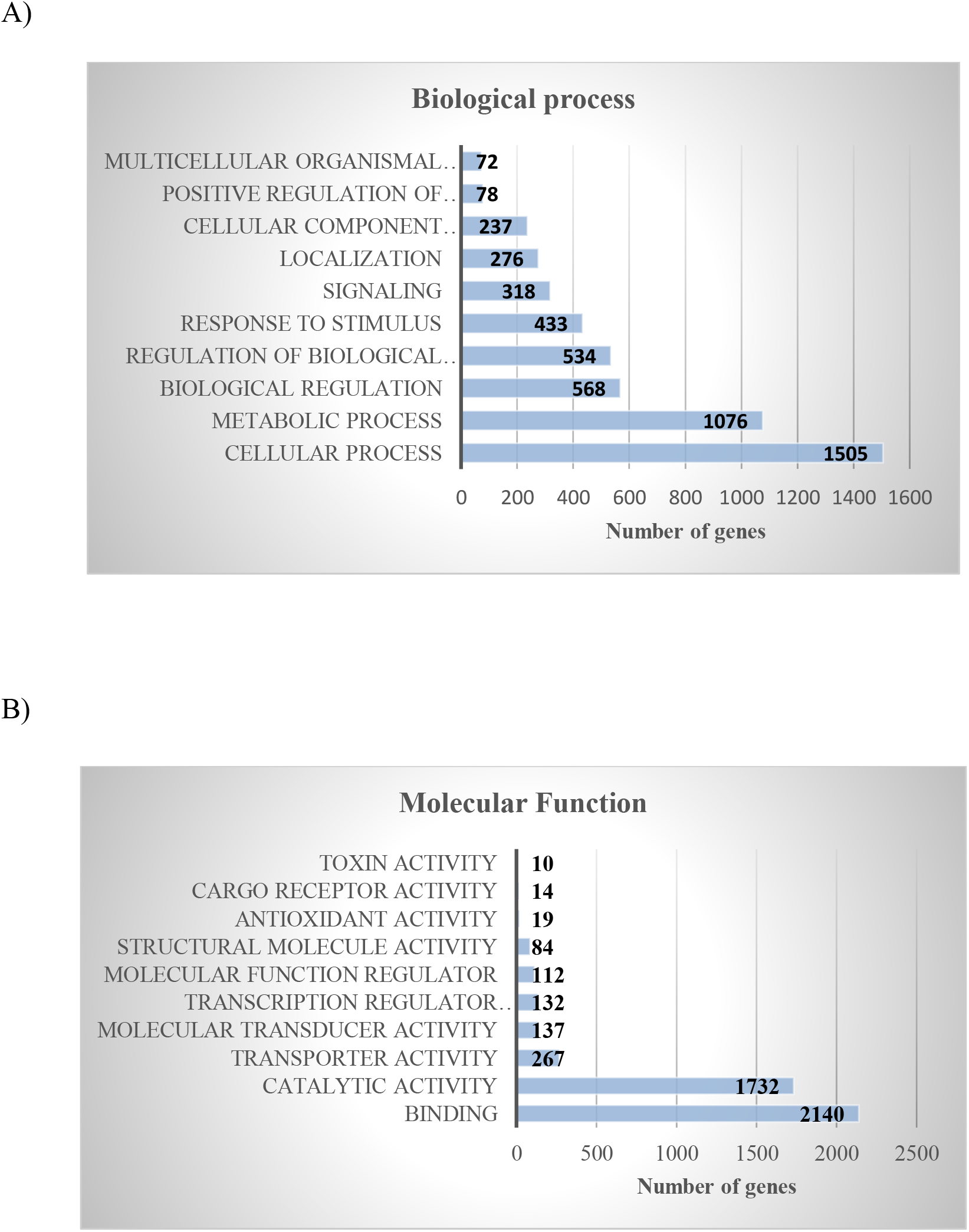
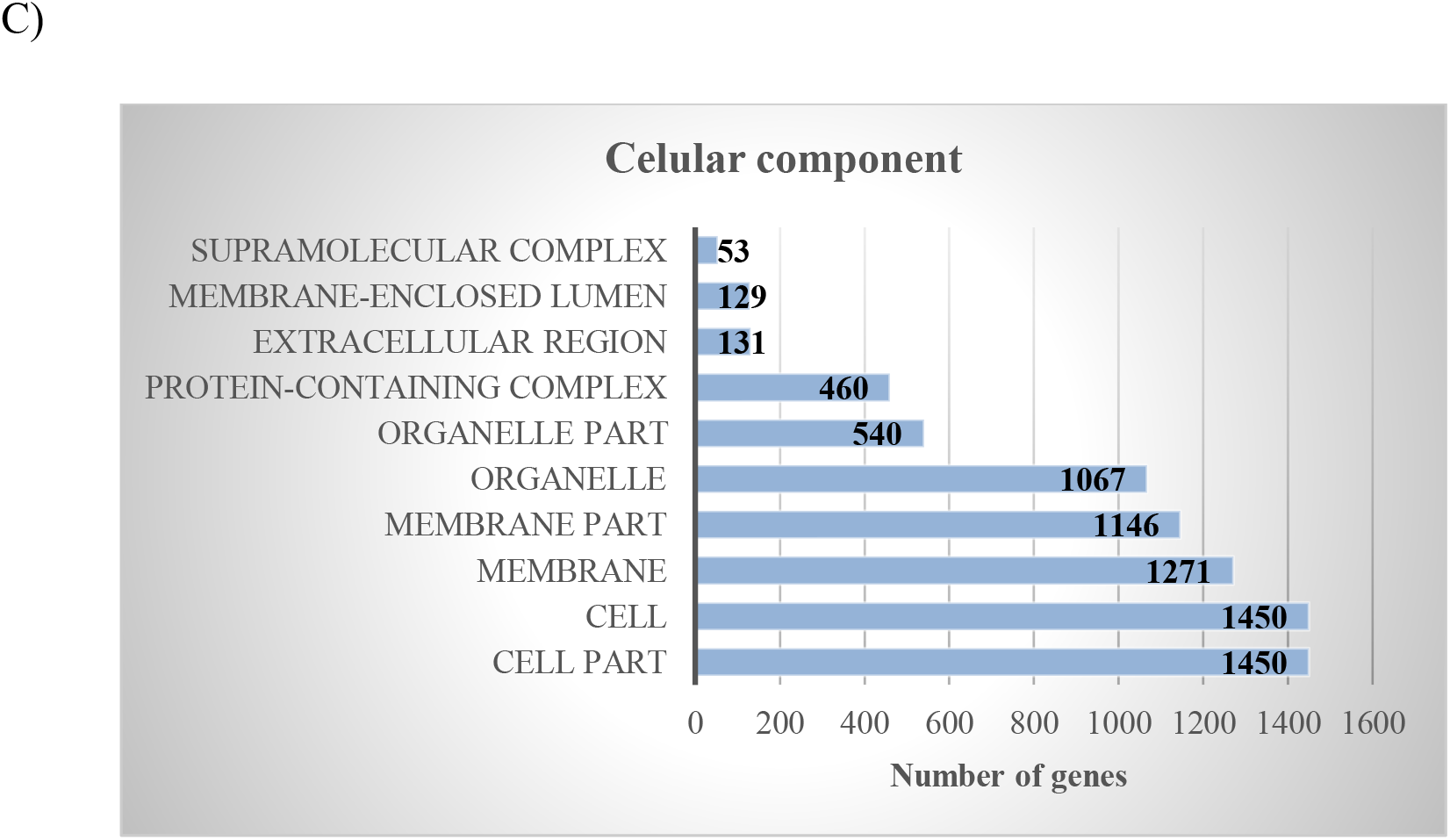
Top 10 gene ontology (GO) terms according to A) biological process B) molecular function, and C) cellular component categories

### Differentially expressed genes (DEGs)

Pairwise comparisons performed across time and between levels of infection yielded a total of 3656 DEGs in the different comparisons performed, some of them up-or down-regulated in more than one comparison (Table S3). The proportion of DEGs with respect to the genes annotated in each chromosome was rather homogeneous, although C6, C16 and C19 were particularly enriched (> 35%) (Figure 6). The maximum amount of DEGs was found in temporal pairwise comparisons involving the digestive gland (T1 *vs* T2: 2009 DEGs; Noia control *vs* T1: 1513 DEGs; Noia control *vs* T2: 1082; Table S4), which suggested that Noia control could be inflating these figures due to the specific environmental or biological factors occurring at the bed of Noia. So, to be conservative, we decided to consider only the set of shared genes between temporal comparisons for this organ, retaining a total of 95 DEGs. Another set of 495 DEGs was detected when comparing the whole meat samples (T0 *vs* T3). We paid special attention to the pairwise comparisons of the infection level within time for the digestive gland (T1 and T2), the target organ of the parasite, taking non-infected individuals as reference (I0). As expected, in both cases the number of DEGs increased with the level of infection (T1: 2, 18, 25, 311 DEGs for I1, I2, I3 and I4, respectively; T2: 6, 68, 87 DEGs for I1, I2 and I3, respectively; Table S4). All in all, a total of 973 DEGs were detected gathering temporal and infection samples according to the criteria exposed (Table S5).

**Figure 6.**
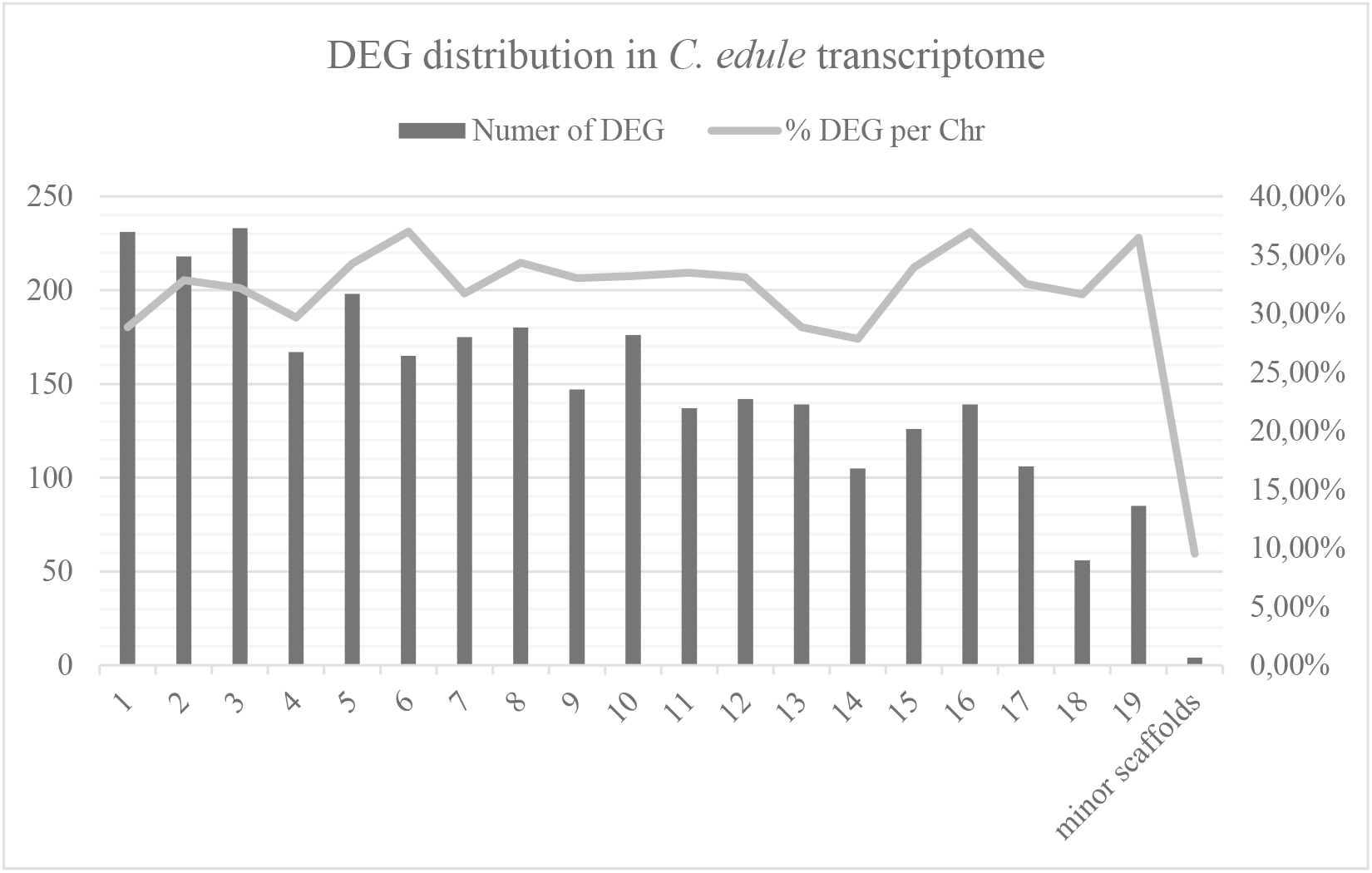
Distribution of differentially expressed genes of *Cerastoderma edule*.in response to *Marteilia cochillia* per chromosome.

### Immune response genes of Cerastoderma edule against marteliosis

Cockles as other bivalve molluscs can thrive in microbe-rich environments as filter-feeders thanks to the strong capability of their innate immune response. Understanding the interaction between the host and the parasite across the infection process is essential to identify key genes involved in the host’s immune response to control this parasitosis. This is the first time where immune-related genes have been massively identified in the common cockle transcriptome and constitutes a step forward towards controlling marteliosis. Table 4 shows some of the relevant immune genes found in our study, which includes, among others, many related to tumour necrosis factor (TNF) and toll-like receptor, but also other key genes in the immune response, such as complement, interferon, interleukin, defensin, membrane-surface binding ligand, ubiquitin, and iron-binding related genes.

**Table 4:** Differentially expressed genes involved in immune response in *Cerastoderma edule* after infection with *Marteilia cochillia*.

| Gene Annotation | Species hit | Gene names |
| --- | --- | --- |
| Putative tumor necrosis factor ligand superfamily member 10-like protein | *Pinctada fucata* | - |
| Lipopolysaccharide and beta-1,3-glucan-binding protein | *Meretrix meretrix* | - |
| Tumor necrosis factor ligand superfamily member | *Crassostrea gigas* | - |
| Peptidoglycan recognition protein 2 | *Mytilus galloprovincialis* | - |
| Tumor Necrosis Factor | *Ruditapes philippinarum* | TNF |
| Toll-like receptor 3 | *Mizuhopecten yessoensis* | KP79_PYT13408 |
| Interferon regulatory factor 2 | *Hyriopsis cumingii* | IRF-2 |
| Tumor necrosis factor ligand superfamily member 11 | *Crassostrea gigas* | CGI_10006440 |
| RNA helicase (EC 3.6.4.13) | *Lingula unguis* | LOC106168940 |
| Tumor necrosis factor | *Ostrea edulis* | - |
| Complement factor B-like protein | *Ruditapes decussatus* | - |
| MyD88 | *Ruditapes philippinarum* | - |
| Polyglutamine-binding protein 1 | *Lingula unguis* | LOC106162394 |
| Complement factor B | *Stylophora pistillata* | Cfb<br>AWC38_SpisGene21788 |
| Interleukin-1 receptor-associated kinase 1-binding protein 1 | *Mizuhopecten yessoensis* | KP79_PYT06648 |
| Toll-like receptor 3 | *Mytilus coruscus* | TLR3 |
| Tumor necrosis factor ligand superfamily member 10 | *Crassostrea gigas* | CGI_10018788 |
| Ferrochelatase (EC 4.99.1.1) | *Salmo salar* | fech |
| RNA helicase (EC 3.6.4.13) | *Mizuhopecten yessoensis* | KP79_PYT17676 |
| Toll-like receptor 3-like | *Neolamprologus brichardi* | - |
| Toll-like receptor f | *Cyclina sinensis* | - |
| TRAF3 | *Sinanodonta woodiana* | TRAF3 |
| E3 ubiquitin-protein ligase TRIM56 (EC 2.3.2.27) | *Bos taurus* | TRIM56 |
| (Tripartite motif-containing protein 56) |  |  |
| Toll-like receptor | *Anthopleura buddemeieri* | TLR |
| Toll-like receptor 1 | *Argopecten irradians* | - |
| Toll-like receptor f | *Mytilus galloprovincialis* | - |
| MYD88 | *Cyclina sinensis* | - |
| Toll-like receptor e | *Mytilus galloprovincialis* | - |
| Toll-like receptor 1 | *Argopecten irradians* | - |
| Poly [ADP-ribose] polymerase (PARP) (EC 2.4.2.-) | *Oryctolagus cuniculus* | PARP14 |
| Tumor necrosis factor ligand superfamily member 10 | *Crassostrea gigas* | CGI_10018788 |
| Hemocyte defensin Cg-Defh1 (Fragment) | *Crassostrea gigas* | 0 |
| Tumor necrosis factor ligand superfamily member 10 | *Mizuhopecten yessoensis* | - |
| Toll-like receptor | *Anthopleura buddemeieri* | TLR |
| Slit-like 1 protein | *Mizuhopecten yessoensis* | KP79_PYT21784 |
| Collectin-12 | *Mesocricetus auratus* | Colec12 |
| Protein toll | *Mizuhopecten yessoensis* | KP79_PYT03261 |
| Toll-like receptor 2 | *Camelus dromedarius* | Cadr_000009830 |
| Protein toll | *Crassostrea gigas* | CGI_10026389 |

### Singular enrichment analysis (SEA)

Singular enrichment analyses were performed considering all DEGs across time and level of infection taking as background the cockle transcriptome here described. Temporal comparisons rendered higher number of enriched functions than comparisons related to the level of infection in accordance with the higher number of genes detected in the former (Table S6). DEGs detected comparing temporal samples involving the cockle whole meat (T0 *vs* T3), were mainly assigned to BP and MF categories and showed enrichment in protein folding biological processes and molecular functions of unfolded protein binding and sulfotransferase activity, while the most reliable comparison involving the digestive gland (T1 *vs* T2) showed a larger list, mainly related to transmembrane transport activities in the MF category and several cilium-related and transport mechanisms in the CC category. These functions may be associated with the digestive gland physiology, which involves intense transport activities and cilium-mediated functions (Lobo-da-Cunha, 2019). DEGs related to comparisons between levels of infection, detected fewer enriched functions as a consequence of the lower number of genes, but associated with immune response and digestive function, such as carbohydrate binding and hydrolase and peptidase activity (Table S6). The former included carbohydrate binding proteins such as lectins which are implicated in host-pathogen interactions and associated with hemocyte membrane (Vasta et al., 2007), while hydrolase activity has been associated with antimicrobial activity in oysters and included some members of the lysozyme family (N-acetylmuramide glycanhydrolase) a member of suggested to be part of the antimicrobial defense mechanism. (McDade and Tripp, 1967).

In summary, this study provides new genomic resources for common cockles, essential to understand their immune response to the parasite *M. cochillia* that has greatly impacted their production in Southern Galician Rias in the last decade. Assembly and characterization of common cockle transcriptome, mostly focused on the digestive gland, has been the basis for identifying DEGs related to response to marteiliosis. This information will be crucial to identify and validate candidate genes and markers for controlling this pathology and recovering cockle’s production.

## Notes

### Competing Interest Statement

The authors have declared no competing interest.

## Literature cited

Allam, B., and Raftos, D. (2015). Immune responses to infectious diseases in bivalves. J. Invertebr. Pathol. 131, 121–136. doi:10.1016/j.jip.2015.05.005.

Altschul, S. F., Gish, W., Miller, W., Myers, E. W., and Lipman, D. J. (1990). Basic local alignment search tool. J. Mol. Biol. 215, 403–410. doi:10.1016/S0022-2836(05)80360-2.

Barcia, R. (1997). The 28S fraction of rRNA in molluscs displays electrophoretic behaviour different from that of mammal cells. Biochem. Mol. Biol. Int. 42, 1089–1092. doi:10.1080/15216549700203551.

Bolger, A. M., Lohse, M., and Usadel, B. (2014). Trimmomatic: A flexible trimmer for Illumina sequence data. Bioinformatics 30, 2114–2120. doi:10.1093/bioinformatics/btu170.

Bray, N. L., Pimentel, H., Melsted, P., and Pachter, L. (2016). Near-optimal probabilistic RNA-seq quantification. Nat. Biotechnol. 34, 525–528. doi:10.1038/nbt.3519.

Bruzos, A. L., Santamarina, M., García-souto, D., Díaz, S., Rocha, S., Zamora, J., et al. (2022). The evolution of two transmissible leukaemias colonizing the coasts of Europe. bioRixiv. doi:10.1101/2022.08.06.503021.

Carrasco, N., Hine, P. M., Durfort, M., Andree, K. B., Malchus, N., Lacuesta, B., et al. (2013). Marteilia cochillia sp. nov., a new *Marteilia* species affecting the edible cockle *Cerastoderma edule* in European waters. Aquaculture 412–413, 223–230. doi:10.1016/j.aquaculture.2013.07.027.

Carrasco, N., Roque, A., Andree, K. B., Rodgers, C., Lacuesta, B., and Furones, M. D. (2011). A Marteilia parasite and digestive epithelial virosis lesions observed during a common edible cockle *Cerastoderma edule* mortality event in the Spanish Mediterranean coast. Aquaculture 321, 197–202. doi:10.1016/j.aquaculture.2011.09.018.

Carss, D. N., Brito, A. C., Chainho, P., Ciutat, A., de Montaudouin, X., Fernández Otero, R. M., et al. (2020). Ecosystem services provided by a non-cultured shellfish species: The common cockle *Cerastoderma edule*. Mar. Environ. Res. 158. doi:10.1016/j.marenvres.2020.104931.

Dairain, A., Maire, O., Meynard, G., Richard, A., Rodolfo-Damiano, T., and Orvain, F. (2020). Sediment stability: can we disentangle the effect of bioturbating species on sediment erodibility from their impact on sediment roughness? Mar. Environ. Res. 162, 105147. doi:10.1016/j.marenvres.2020.105147.

de la Ballina, N. R., Villalba, A., and Cao, A. (2018). Proteomic profile of the European flat oyster *Ostrea edulis* haemolymph in response to bonamiosis and identification of candidate proteins as markers of resistance. Dis. Aquat. Organ. 128, 127–145. doi:10.3354/dao03220.

De Lorgeril, J., Petton, B., Lucasson, A., Perez, V., Stenger, P. L., Dégremont, L., et al. (2020). Differential basal expression of immune genes confers *Crassostrea gigas* resistance to Pacific oyster mortality syndrome. BMC Genomics 21, 63. doi:10.1186/s12864-020-6471-x.

de Montaudouin, X. de, Arzul, I., Cao, A., Carballal, M. J., Chollet, B., Correia, S., et al. (2021). Catalogue of Parasites and Diseases of the Common Cockle Cerastoderma edule - COCKLES PROJECT. UA Editora – Universidade de Aveiro doi:10.34624/9a9c-9j21.

Dobin, A., Davis, C. A., Schlesinger, F., Drenkow, J., Zaleski, C., Jha, S., et al. (2013). STAR: Ultrafast universal RNA-seq aligner. Bioinformatics 29, 15–21. doi:10.1093/bioinformatics/bts635.

Ducrotoy, J.-P., Rybarczyk, H., Souprayen, S., Bachelet, G., Beukema, J. J., Desprez, M., et al. (1991). A comparison of the population dynamics of the cockle (Cerastoderma edule, L.), in North-Western Europe. University of Caen, France: Proceedings of the Estuarine and Coastal Sciences; ECSA 19; 4–8 September 1989. University of Caen, France.

FAO (2021). Food and Agriculture Organization of the United Nations. Statistics. Available at: http://www.fao.org/fishery/statistics/global-production/en [Accessed June 29, 2021].

Farhat, S., Tanguy, A., Pales, E., Guo, X., Boutet, I., Smolowitz, R., et al. (2020). Identification of variants associated with hard clam, *Mercenaria mercenaria*, resistance to Quahog Parasite Unknown disease. Genomics 112, 4887–4896. doi:10.1016/j.ygeno.2020.08.036.

Fernández-Boo, S., Villalba, A., and Cao, A. (2016). Protein expression profiling in haemocytes and plasma of the Manila clam *Ruditapes philippinarum* in response to infection with Perkinsus olseni. J. Fish Dis. 39, 1369–1385. doi:10.1111/jfd.12470.

Gómez-Chiarri, M., Warren, W. C., Guo, X., and Proestou, D. (2015). Developing tools for the study of molluscan immunity: The sequencing of the genome of the eastern oyster, *Crassostrea virginica*. Fish Shellfish Immunol. 46, 2–4. doi:10.1016/j.fsi.2015.05.004.

Gutierrez, A. P., Bean, T. P., Hooper, C., Stenton, C. A., Sanders, M. B., Paley, R. K., et al. (2018). A genome-wide association study for host resistance to ostreid herpesvirus in Pacific oysters *(Crassostrea gigas)*. G3 Genes, Genomes, Genet. 8, 1273–1280. doi:10.1534/g3.118.200113.

Gutierrez, A. P., Symonds, J., King, N., Steiner, K., Bean, T. P., and Houston, R. D. (2020). Potential of genomic selection for improvement of resistance to ostreid herpesvirus in Pacific oyster *(Crassostrea gigas)*. Anim. Genet. 51, 249–257. doi:10.1111/age.12909.

Hasanuzzaman, A. F. M., Cao, A., Ronza, P., Fernández-Boo, S., Rubiolo, J. A., Robledo, D., et al. (2020). New insights into the Manila clam – *Perkinsus olseni* interaction based on gene expression analysis of clam hemocytes and parasite trophozoites through in vitro challenges. Int. J. Parasitol. 50, 195–208. doi:10.1016/j.ijpara.2019.11.008.

He, Y., Yu, H., Bao, Z., Zhang, Q., and Guo, X. (2012). Mutation in promoter region of a serine protease inhibitor confers *Perkinsus marinus* resistance in the eastern oyster (*Crassostrea virginica*). Fish Shellfish Immunol. 33, 411–417. doi:10.1016/j.fsi.2012.05.028.

Houston, R. D., Bean, T. P., Macqueen, D. J., Gundappa, M. K., Jin, Y. H., Jenkins, T. L., et al. (2020). Harnessing genomics to fast-track genetic improvement in aquaculture. Nat. Rev. Genet. 21, 389–409. doi:10.1038/s41576-020-0227-y.

Howard, D. W., Lewis, E. J., Keller, B. J., and Smith, C. S. (2004). Histological techniques for marine bivalve mollusks and crustaceans. NOOA Tech. Memo. NOS NCCOS 5, 218 pp. Available at: http://hdl.handle.net/1834/30812.

Iglesias, D., Villalba, A., Cao, A., and Carballal, M.. (2019). Is natural selection enhancing resistance against marteiliosis in cockles recruited in the inner side of the ria of Arousa? in 19th International Conference on Diseases of Fish and Shellfish, Oporto, Portugal (Porto, Portugal: European Association of Fish Pathologists), pp.184. Available at: https://eafp.org/wp-content/uploads/2020/01/2019-porto-abstract-book.pdf.

Iglesias, D., Villalba, A., Darriba, S., Cao, A., Mariño, C., Fernández, J., et al. (2017). Epidemiological patterns of marteiliosis affecting the common cockle *Cerastoderma edule* in Galicia (NW Spain). in Book of Abstracts:18th International Conference on Diseases of Fish and Shellfish. (Belfast (UK): European Association of Fish Pathologists), pp.215. Available at: https://eafp.org/wp-content/uploads/2018/10/18-eafp-abstract-book.pdf.

Iglesias, D., Villalba, A., Darriba, S., Mariño, C., Fernández, J., and Carballal, M. J. (2015). Cockle *Cerastoderma edule* marteliosis first detected in Ría de Arousa (Galicia, Nw Spain) has spread to other Galician rias causing mass mortality. in 17th International Conference on Diseases of fish and Shellfish (Las Palmas de Gran canaria: European Association of Fish Pathologists), 348.

Iglesias, D., Villalba, A., Mariño, C., No, E., and Carballal, M.. Long-term survey discloses a shift in the dynamics pattern of a lethal emerging marine disease and raises hypothesis to explain it. The case of marteiliosis of cockles *Cerastoderma edule*. Mar. Biol.

Jackson-Bué, M., Brito, A. C., Cabral, S., Carss, D. N., Carvalho, F., Chainho, P., et al. (2022). Inter-country differences in the cultural ecosystem services provided by cockles. People Nat. 4, 71–87. doi:10.1002/pan3.10252.

Kelley, K. N. (2009). Use of Recycled Oyster Shells as Aggregate for Previous Concrete. MSC Thesis. Available at: http://ufdcimages.uflib.ufl.edu/UF/E0/04/12/69/00001/kelley_k.pdf.

La Peyre, J. F., Casas, S. M., Richards, M., Xu, W., and Xue, Q. (2019). Testing plasma subtilisin inhibitory activity as a selective marker for dermo resistance in eastern oysters. Dis. Aquat. Organ. 133, 127–139. doi:10.3354/dao03344.

Leprêtre, M., Faury, N., Segarra, A., Claverol, S., Degremont, L., Palos-ladeiro, M., et al. (2021). Comparative Proteomics of Ostreid Herpesvirus 1 and Pacific Oyster Interactions With Two Families Exhibiting Contrasted Susceptibility to Viral Infection. Front. Immunol. 11, 621994. doi:10.3389/fimmu.2020.621994.

Li, H., Handsaker, B., Wysoker, A., Fennell, T., Ruan, J., Homer, N., et al. (2009). The Sequence Alignment/Map format and SAMtools. Bioinformatics 25, 2078–2079. doi:10.1093/bioinformatics/btp352.

Longshaw, M., and Malham, S. K. (2013). A review of the infectious agents, parasites, pathogens and commensals of European cockles *(Cerastoderma edule* and *C. glaucum)*. J. Mar. Biol. Assoc. United Kingdom 93, 227–247. doi:10.1017/s0025315412000537.

Malham, S. K., Hutchinson, T. H., and Longshaw, M. (2012). A review of the biology of European cockles (Cerastoderma spp.). J. Mar. Biol. Assoc. United Kingdom 92, 1563–1577. doi:10.1017/S0025315412000355.

McDade, J. E., and Tripp, M. R. (1967). Lysozyme in oyster mantle mucus. J. Invertebr. Pathol. 9, 581–582. doi:10.1016/0022-2011(67)90146-2.

Meistertzheim, A. L., Calvès, I., Rousse, V., Van Wormhoudt, A., Laroche, J., Huchette, S., et al. (2014). New genetic markers to identify European resistant abalone to vibriosis revealed by high-resolution melting analysis, a sensitive and fast approach. Mar. Biol. 161, 1883–1893. doi:10.1007/s00227-014-2470-2.

Morris, J. P., Backeljau, T., and Chapelle, G. (2018). Shells from aquaculture: a valuable biomaterial, not a nuisance waste product. Rev. Aquac. 11, 42–57. doi:10.1111/raq.12225.

Nagalakshmi, U., Wang, Z., Waern, K., Shou, C., Raha, D., Gerstein, M., et al. (2008). The Transcriptional Landscape of the Yeast Genome Defined by RNA Sequencing. Science (80-.). 320, 1344–1349. doi:10.1126/science.1158441.

Nie, Q., Yue, X., and Liu, B. (2015). Development of Vibrio spp. infection resistance related SNP markers using multiplex SNaPshot genotyping method in the clam *Meretrix meretrix*. Fish Shellfish Immunol. 43, 469–476. doi:10.1016/j.fsi.2015.01.030.

Nikapitiya, C., McDowell, I. C., Villamil, L., Muñoz, P., Sohn, S. B., and Gomez-Chiarri, M. (2014). Identification of potential general markers of disease resistance in American oysters, *Crassostrea virginica* through gene expression studies. Fish Shellfish Immunol. 41, 27–36. doi:10.1016/j.fsi.2014.06.015.

Pertea, M., Pertea, G. M., Antonescu, C. M., Chang, T. C., Mendell, J. T., and Salzberg, S. L. (2015). StringTie enables improved reconstruction of a transcriptome from RNA-seq reads. Nat. Biotechnol. 33, 290–295. doi:10.1038/nbt.3122.

Pimentel, H., Bray, N. L., Puente, S., Melsted, P., and Pachter, L. (2017). Differential analysis of RNA-seq incorporating quantification uncertainty. Nat. Methods 14, 687–690. doi:10.1038/nmeth.4324.

Potts, R. W. A., Gutierrez, A. P., Penaloza, C. S., Regan, T., Bean, T. P., and Houston, R. D. (2021). Potential of genomic technologies to improve disease resistance in molluscan aquaculture. Philos. Trans. R. Soc. B Biol. Sci. 376, 20200168. doi:10.1098/rstb.2020.0168.

Proestou, D. A., and Sullivan, M. E. (2020). Fish and Shellfish immunology variation in global transcriptomic response to *Perkinsus marinus* infection among eastern oyster families highlights potential mechanisms of disease resistance. Fish Shellfish Immunol. 96, 141–151. doi:10.1016/j.fsi.2019.12.001.

Sambade, I. M., Casanova, A., Blanco, A., Gundappa, M. K., Bean, T. P., Macqueen, D. J., et al. (2022). A single genomic region involving a putative chromosome rearrangement in flat oyster (*Ostrea edulis*) is associated with differential host resilience to the parasite *Bonamia ostreae*. Evol. Appl. 15, 1408–1422. doi:10.1111/eva.13446.

Schultz, J. H., and Adema, C. M. (2017). Comparative immunogenomics of molluscs. Dev. Comp. Immunol. 75, 3–15. doi:10.1016/j.dci.2017.03.013.

Simonian, M., Nair, S. V., O’Connor, W. A., and Raftos, D. A. (2009). Protein markers of *Marteilia sydneyi* infection in Sydney rock oysters, *Saccostrea glomerata*. J. Fish Dis. 32, 367–375. doi:10.1111/j.1365-2761.2009.01022.x.

Smits, M., Enez, F., Ferraresso, S., Dalla Rovere, G., Vetois, E., Auvray, J. F., et al. (2020). Potential for Genetic Improvement of Resistance to *Perkinsus olseni* in the Manila Clam, *Ruditapes philippinarum*, Using DNA Parentage Assignment and Mass Spawning. Front. Vet. Sci. 7, 579840. doi:10.3389/fvets.2020.579840.

Tebble, N. (1976). British Bivalve Seashells: A Handbook for Identification. Royal Scottish Museum Available at: https://books.google.es/books?id=3MwgAQAAIAAJ.

Thieltges, D. W. (2006). Parasite induced summer mortality in the cockle Cerastoderma edule by the trematode *Gymnophallus choledochus*. Hydrobiologia 559, 455–461. doi:10.1007/s10750-005-1345-4.

Tian, T., Liu, Y., Yan, H., You, Q., Yi, X., Du, Z., et al. (2017). AgriGO v2.0: A GO analysis toolkit for the agricultural community, 2017 update. Nucleic Acids Res. 45, W122–W129. doi:10.1093/nar/gkx382.

Tyler-Walters, H. (2007). *Cerastoderma edule* (Common cockle). Tyler-Walters H. Hiscock K. Mar. Life Inf. Netw. Biol. Sensit. Key Inf. Rev. [on-line]. Plymouth Mar. Biol. Assoc. United Kingdom. doi:10.17031/marlinsp.1384.1.

Vaibhav, V., Thompson, E. L., Raftos, D. A., and Haynes, P. A. (2018). Potential protein biomarkers of QX disease resistance in selectively bred Sydney Rock Oysters. Aquaculture 495, 144–152. doi:10.1016/j.aquaculture.2018.05.035.

van der Schatte Olivier, A., Jones, L., Vay, L. Le, Christie, M., Wilson, J., and Malham, S. K. (2020). A global review of the ecosystem services provided by bivalve aquaculture. Rev. Aquac. 12, 3–25. doi:10.1111/raq.12301.

Vasta, G. R., Ahmed, H., Tasumi, S., Odom, E. W., and Saito, K. (2007). Biological Roles of Lectins in Innate Immunity: Molecular and Structural Basis for Diversity in Self/Non-Self Recognition. in Current Topics in Innate Immunity, ed. J. D. Lambris (New York, NY: Springer New York), 389–406. doi:10.1007/978-0-387-71767-8_27.

Vera, M., Pardo, B. G., Cao, A., Vilas, R., Fernández, C., Blanco, A., et al. (2019). Signatures of selection for bonamiosis resistance in European flat oyster (*Ostrea edulis*. *New genomic tools for breeding programs and management of natural resources*. Evol. Appl. 12, 1781–1796. doi:10.1111/eva.12832.

Villalba, A., Iglesias, D., Ramilo, A., Darriba, S., Parada, J. M., No, E., et al. (2014). Cockle *Cerastoderma edule* fishery collapse in the Ría de Arousa (Galicia, NW Spain) associated with the protistan parasite *Marteilia cochillia*. Dis. Aquat. Organ. 109, 55–80. doi:10.3354/dao02723.

Wang, K., del Castillo, C., Corre, E., Espinosa, E. P., and Allam, B. (2016). Clam focal and systemic immune responses to QPX infection revealed by RNA-seq technology. BMC Genomics 17, 146. doi:10.1186/s12864-016-2493-9.

Wang, W., Song, X., Wang, L., and Song, L. (2018). Pathogen-derived carbohydrate recognition in molluscs immune defense. Int. J. Mol. Sci. 19, 721. doi:10.3390/ijms19030721.

Watson, A., Agius, J., Ackerly, D., Beddoe, T., and Helbig, K. (2022). The Role of Anti-Viral Effector Molecules in Mollusc Hemolymph. Biomolecules 12, 345. doi:10.3390/biom12030345.

Wolff, W. J. (2005). The exploitation of living resources in the Dutch Wadden Sea: A historical overview. Helgol. Mar. Res. 59, 31–38. doi:10.1007/s10152-004-0204-4.

Ye, J., Fang, L., Zheng, H., Zhang, Y., Chen, J., Zhang, Z., et al. (2006). WEGO: A web tool for plotting GO annotations. Nucleic Acids Res. 34, 293–297. doi:10.1093/nar/gkl031.

Ye, J., Zhang, Y., Cui, H., Liu, J., Wu, Y., Cheng, Y., et al. (2018). WEGO 2.0: A web tool for analyzing and plotting GO annotations, 2018 update. Nucleic Acids Res. 46, W71–W75. doi:10.1093/nar/gky400.

